# Disentangling the cascading effects of Grapevine Red Blotch Virus infection on vine physiology

**DOI:** 10.1101/2023.06.15.545163

**Authors:** Cody R. Copp, Joseph B. DeShields, Suraj Kar, Claire Kirk, Ricky Clark, Marianna Stowasser, Achala N. KC, Alexander D. Levin

## Abstract

Grapevine Red Blotch Virus is a major grapevine pathogen and is associated with reduced carbon assimilation and delayed berry ripening in *Vitis vinifera* L. Recent work suggests that the virus alters leaf carbon metabolism prior to emergence of visible symptoms. Therefore, diurnal and seasonal measurements were conducted to quantify changes in leaf carbon balance and to elucidate the chronology of symptom progression in leaves and fruit. Healthy and infected vines were assayed in a commercial vineyard during which leaf water relations, photosynthesis, and nonstructural carbohydrates were measured. Additionally, sugar and anthocyanin accumulation in the fruit were monitored at the end of the season to characterize the impact of the virus on ripening. Virus infection significantly reduced carbon assimilation pre- and postveraison, but the impact was more pronounced postveraison and during the afternoon when vine water status was the lowest. Similarly, virus infection significantly increased leaf starch concentration pre- and postveraison, but increased leaf starch in infected vines was detected two weeks prior to veraison. Virus infection had the greatest impact on obstructing leaf carbon export postveraison, especially during the afternoon. The virus had no impact on chlorophyll fluorescence, indicating there was no sustained photosystem impairment and suggesting that changes in chlorophyll fluorescence were a transient response to reduced carbon assimilation and export. This study provides evidence that reduced carbon export constitutes a feedback inhibition response to accumulation of leaf starch prior to the appearance of visible symptoms or impacts to ripening, which may aid earlier detection of the virus.

---

Grapevine Red Blotch Virus (GRBV) is a phloem-limited monopartite geminivirus of grapevine (*Vitis vinifera* L.) in the family *Geminiviridae* (Sudarshana et al., 2015), and is the causal agent of Grapevine Red Blotch Disease (GRBD) (Yepes et al., 2018). GRBD was first documented in California vineyards in 2008 (Calvi, 2011) and GRBV has subsequently been found in diverse viticultural regions around the world (Rumbaugh et al., 2021). GRBD significantly reduces fruit quality primarily by impairing sugar accumulation in the berry and, in case of red-fruited cultivars, by reducing skin anthocyanin accumulation (Girardello et al., 2019; Martínez-Lüscher et al., 2019; Levin and KC, 2020). With global wine trade value recently estimated at nearly $35 billion (OIV, 2022), it is crucial to understand the physiological impacts of this disease on grapevines to improve disease management strategies.

GRBV has been identified in archival plant material from the 1940s (Al Rwahnih et al., 2015). It is primarily spread through propagation of infected plant material, but it can also be less efficiently vectored by insects in the *Membracidae* family (Bahder et al., 2016; Dalton et al., 2019). Nevertheless, rapid disease progression through vineyards can be as large as 30-fold over three years (KC et al., 2022). The primary visible symptoms of viral infection in the plant consist of the eponymous foliar reddening (red blotches) and are accompanied by impaired carbon assimilation (*A*_net_) and fruit ripening (Martínez-Lüscher et al., 2019; Levin and KC, 2020). Blanco-Ulate et al. (2017) reported that GRBV impacts ripening by hindering transcriptional regulation of anthocyanin biosynthesis and by impairing hormonal signaling pathways for abscisic acid, ethylene, and auxin. Martínez-Lüscher et al. (2019) measured increased concentrations of foliar nonstructural carbohydrates (NSC) together with lower total soluble solids at harvest and concluded that the reduction in berry sugar accumulation was due to impaired sugar translocation from source to sink.

Increased foliar NSC concomitant with increased foliar anthocyanins suggests feedback inhibition of *A*_net_ whereby excess NSC is a signal to downregulate *A*_net_ and also a substrate for biosynthesis of photoprotective flavanoids (Lee and Gould, 2002). Such a mechanism has been implicated in other grapevine-virus pathosystems, like Grapevine leafroll virus (Halldorson and Keller, 2018), and with other phloem-limited pathogens of woody perennial crops such as Huanglongbing (HLB) in citrus (Etxeberria et al. 2009). Girdling experiments have shown that mechanical injury to the phloem causes an accumulation of foliar NSC and subsequent reduction of *A*_net_ (Roper and Williams, 1989). In response to biotic stress from pathogens, host-defense responses, such as callose formation around plasmodesmata, are triggered in the phloem to prevent systemic infection (Lewis et al., 2022), and could also cause an excess accumulation of foliar NSC. It has been further suggested that reduced *A*_net_ and translocation rates contribute to other reported GRBD symptoms such as reduced stomatal conductance (*g*_sw_) and increased stem water potential (Ψ_stem_) (Martínez-Lüscher et al., 2019; Levin and KC, 2020). However, these associations have only been inferred from seasonal changes to *A*_net_ and limited foliar NSC measurements late in the season, without quantification of the hypothesized inhibition of sugar translocation.

Recent studies on red oak (*Quercus rubra*) and grapevine presented a mass-balance approach to estimating carbon export that could be useful in a GRBV context to quantify inhibition of sugar translocation (Gersony et al., 2020; Dayer et al., 2021). Evidence from healthy vines suggests that there is diurnal variation in both the makeup and synthesis of foliar NSC, a disruption of which could be an indication of the impact of GRBV on carbohydrate production and export throughout the day (Chaumont et al., 1994; Yu et al., 2009). Dayer et al. (2021) and Gersony et al. (2020) integrated these parameters to show that there is considerable diurnal variation in carbon export rate in grapevine and red oak, respectively.

Chlorophyll fluorescence (ChlF), an interconnected parameter with *A*_net_, can provide another metric to disentangle the impact of GRBV on photosynthetic machinery. In tandem with *A*_net_, ChlF parameters exhibit normal diurnal variation (Ding et al., 2006) that could be altered under GRBV infection. Synthesis of photoprotective flavonoids such as anthocyanin on infected leaves could be an indicator of an effect of GRBV on photosystem II (Feild et al., 2001). Additionally, specific ChlF parameters like maximum photochemical efficiency of photosystem II (*F*_v_/*F*_m_) indicate whether there is sustained damage to the photosystem that is often attributed to biotic or irreversible abiotic damage (Maxwell and Johnson, 2000; Gallé and Flexas, 2010).

The present study was conducted to quantify changes in leaf carbon balance and test the hypothesis that GRBV reduces leaf carbon export. Additionally, we hypothesized that there is diurnal variation with respect to the impact of GRBV on leaf carbon metabolism and chlorophyll fluorescence. Field measurements on healthy and infected vines were conducted during which leaf water potential (Ψ_leaf_), leaf gas exchange, and ChlF were quantified, and those same leaves were collected for NSC analysis. Leaf carbon export was estimated using a mass balance approach involving changes in *A*_net_ and NSC. Sugar and anthocyanin accumulation in the fruit was monitored weekly to characterize the impact of GRBV throughout ripening. This study is aimed at further disentangling the impacts of GRBV on vine physiology and fruit ripening while potentially increasing the ability to detect changes within the vine before the appearance of red blotches to aid in disease management.

## RESULTS

### Environmental conditions

Environmental conditions during the diurnal measurements in 2020 were observed to be normal for the growing region, however, air temperature (T_air_) and VPD were higher compared to the historical average during the postveraison sample date (Figs. 1A-D). Preveraison, T_air_ increased steadily from 1400 hr. until a maximum of approximately 31 °C was recorded at 1800 hr. Postveraison, the maximum T_air_ of 38 °C was recorded at 1600 hr. Following a similar trend, VPD reached maximum values during the same time periods as T_air_, with values peaking at 3.8 and 5.8 kPa for preveraison and postveraison, respectively.

**Figure 1.**
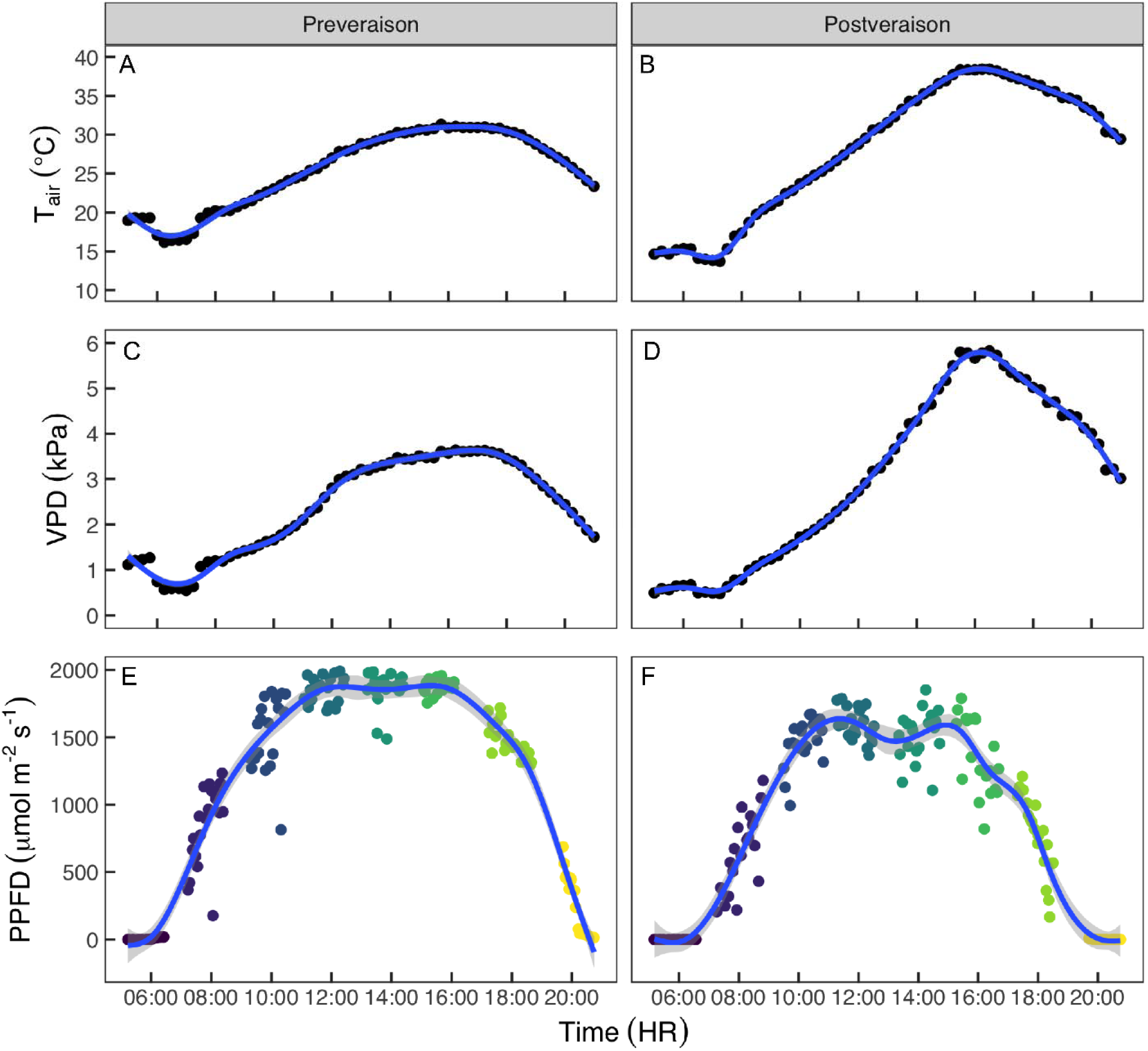
Diurnal courses of ambient air temperature (T_air_; A, B), vapor pressure deficit (VPD; C, D), and photosynthetic photon flux density (PPFD; E, F) at the study site on 23 July (preveraison; A, C, E) and 3 September (postveraison; B, D, F). T_air_ and VPD data were 15-min averages recorded by the nearby weather station, while PPFD data were recorded by the LI-6400XT sensor head during photosynthesis measurements. Colors represent data collected during one sampling round. Smoothed lines show a 1-hr. running average ± 95% confidence intervals.

Saturating PPFD (>1500 µmol m^-2^ s^-1^; Smart, 1974) was maintained between 1000 and 1600 hr. for both sampling periods (Figs. 1E-F), however, this period was slightly shorter postveraison, as typical for early September compared to late July.

### Leaf water potential and gas exchange

There were no significant effects of virus status on Ψ_leaf_ at any given time of the day, except for the measurement taken at 2000 hr. for preveraison sampling, at which point RB+ vines had a higher Ψ_leaf_ compared to RB-vines (Fig. 2A). At both sampling dates, Ψ_leaf_ fell steadily throughout the course of the day and hit a minimum value between 1500 and 1700 hr. Despite similar predawn values for Ψ_leaf_ between the two sampling dates, the minimum values were lower postveraison (-0.99 MPa) compared to preveraison (-0.73 MPa).

**Figure 2.**
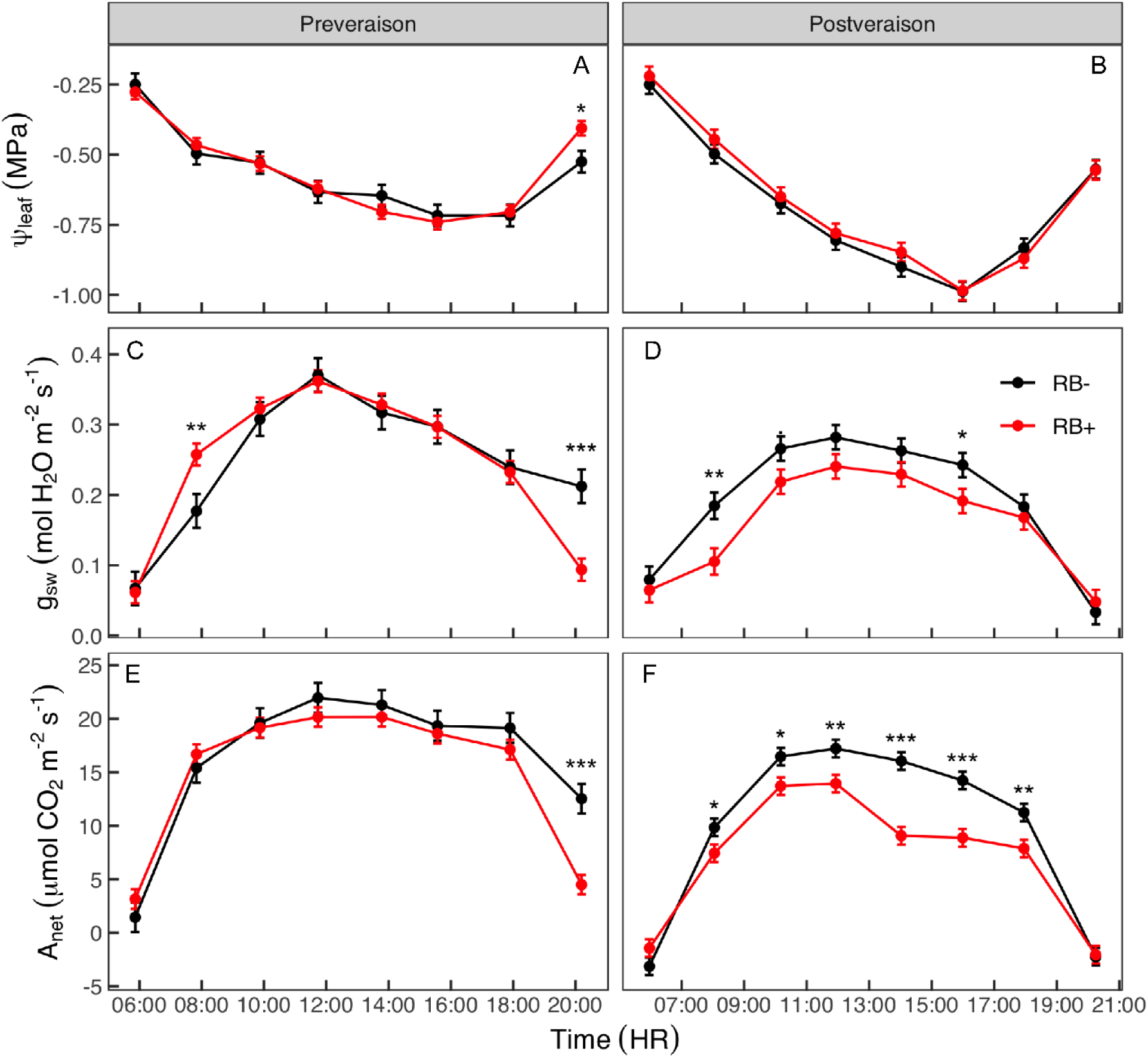
Responses of leaf water potential (Ψ_leaf_; A, B), stomatal conductance to water vapor (*g*_sw_; C, D), and net carbon assimilation (*A*_net_; E, F) between healthy (RB-) and infected (RB+) vines over the course of the day on 23 July (preveraison; A, C, E) and 3 September (postveraison; B, D, F). Data are means ± SE (n = 3-7). ‘***,’ ‘**,’ and ‘*’ represent statistically significant differences between means at *p* < 0.001, 0.01, and 0.05, respectively.

Virus status had a notable impact on gas exchange both in terms of *g*_sw_ and *A*_net_, especially during the postveraison sampling period (Figs. 2D and 2F). During this period RB+ vines showed consistently lower *g*_sw_ and *A*_net_ values throughout the day. Notably, the decreased postveraison *A*_net_ in RB+ vines were more pronounced during the afternoon compared to early morning. Though there were statistically significant interactions among virus status and measurement time for both gas exchange parameters during both sampling dates, significant differences between individual means were limited to early morning and late evening during the preveraison sampling date. When averaged across all measurement times during which light levels were saturating (between 1000 hr. to 1600 hr.), *g*_sw_ and *A*_net_ responses were not significantly different between RB+ and RB-vines preveraison (*p* = 0.773 and 0.229, for *g*_sw_ and *A*_net_, respectively), however, they were significantly different postveraison (*p* = 0.033 and < 0.0001, for *g*_sw_ and *A*_net_, respectively) (data not shown).

### Chlorophyll fluorescence

There were no significant differences in *F*_v_/*F*_m_ between RB+ and RB-vines during either sampling date (Table 1). Averaged across virus status, *F*_v_/*F*_m_ was significantly higher preveraison compared to postveraison (*p* = 0.011), however all values ranged between 0.75 and 0.85 (Table 1). Virus status had notable impact on light-acclimated fluorescence parameters such as Φ_PSII_, *J*, and *qP* during the postveraison sampling date (Figs. 3A-F). All these parameters measured during midday (between 1200 and 1600 hr.) of the postveraison sampling date showed significantly lower values in RB+ vines compared to RB-vines. The impact of GRBV on Φ_PSII_, *J*, and *qP* followed a similar trend as for *A*_net_ whereby there were limited reductions in RB+ vines preveraison but pronounced reductions in RB+ vines postveraison (Figs. 2E-F and 3). Significant reductions in preveraison responses were observed at the end of the day (e.g., increasing reduction in *J* in late afternoon and early evening). When averaged across all measurement times during which light levels were saturating (1000 to 1600 hr.), Φ_PSII_, *J*, and *qP* responses were not significantly different between RB+ and RB-vines during preveraison sampling date (*p* = 0.479, 0.476, and 0.644 for Φ_PSII_, *J*, and *qP,* respectively) (data not shown). However, they were significantly different during postveraison sampling date (*p* = 0.008, < 0.0001, and 0.005, for Φ_PSII_, *J*, and *qP*, respectively) (data not shown). NPQ was higher by an average of 15% during the postveraison sampling day in RB+ vines compared to RB-vines, but it was not statistically significant (Fig. 3H).

**Table 1.** Response of maximum quantum efficiency of dark-acclimated leaves (*F*_v_/*F*_m_) between healthy (RB-) and infected (RB+) vines taken predawn on 23 July (preveraison) and 3 September (postveraison). Data are means ± one standard error (n = 3-7). ANOVA results are shown below data.

| Virus status | $F_v/F_m$ | |
| --- | --- | --- |
|  | Preveraison | Postveraison |
| RB- | $0.807 \pm 0.008$ | $0.795 \pm 0.006$ |
| RB+ | $0.809 \pm 0.005$ | $0.785 \pm 0.006$ |
| Mean | $0.808 \pm 0.005$ | $0.790 \pm 0.004$ |
|  | ANOVA | *p-value* |
|  | Virus | 0.523 |
|  | Date | 0.011 |
|  | V * D | 0.275 |

**Figure 3.**
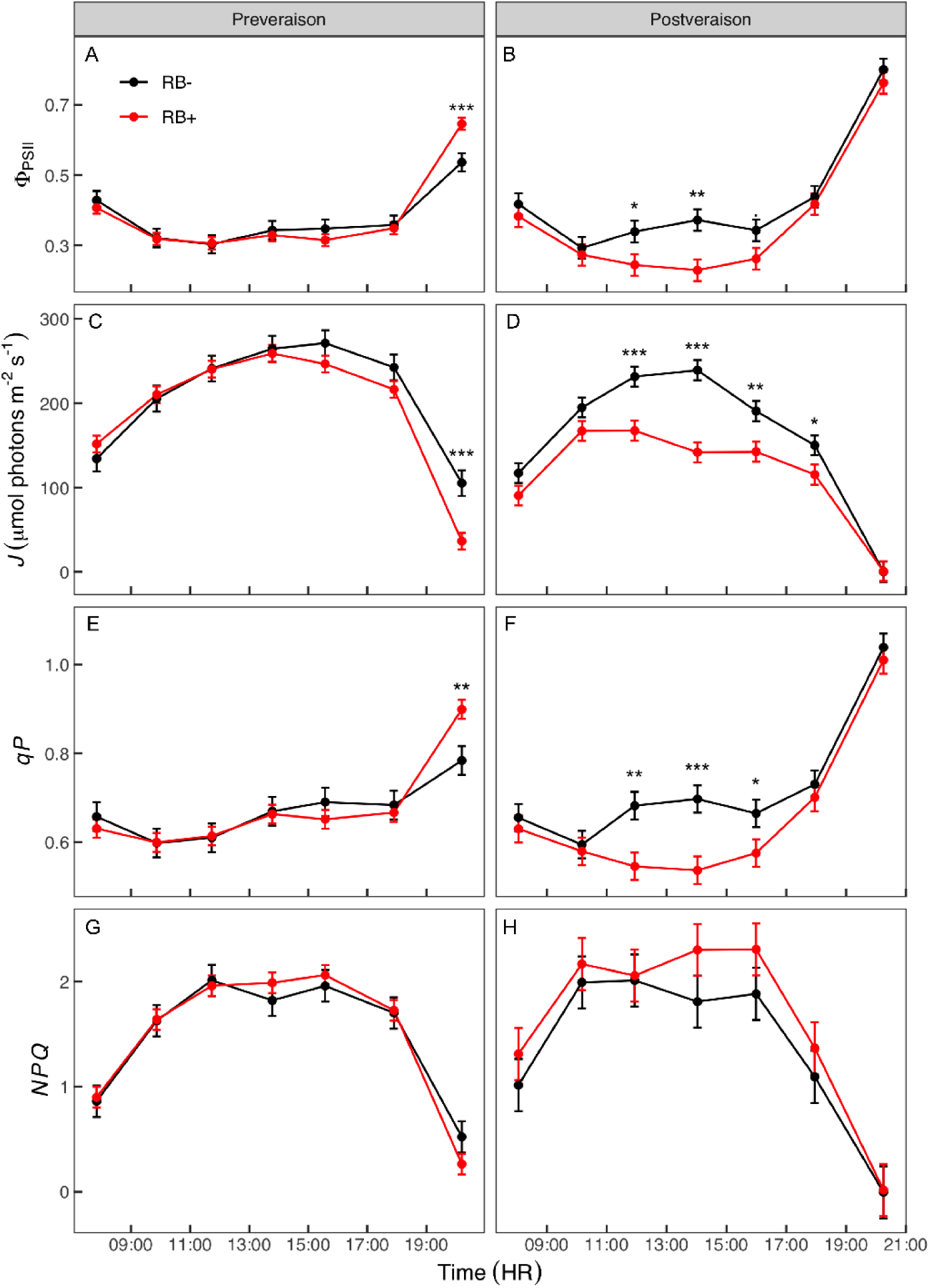
Responses of quantum yield of PSII (Φ_PSII_; A, B), electron transport rate (*J*; C, D), proportion of open PSII (*qP*; E, F), and non-photochemical quenching (*NPQ*; G, H) between healthy (RB-) and infected (RB+) vines over the course of the day on 23 July (preveraison; A, C, E, G) and 3 September (postveraison; B, D, F, H). Data are means ± one standard error (n = 3-7). ‘***,’ ‘**,’ ‘*,’ and ‘.’ represent statistically significant differences between means at *p* < 0.001, 0.01, 0.05, and 0.10, respectively.

### Nonstructural carbohydrates

No differences were found in soluble sugar concentrations between RB+ and RB-leaves during either pre- or postveraison sampling dates (Figs. 4A-B). Foliar soluble sugar concentration followed a significant cubic trend (*p* < 0.05), declining from predawn to mid-morning (∼1000 hr.), followed by an increase to a peak in the afternoon (between 1400 and 1500 hr.) before a late afternoon drop.

**Figure 4.**
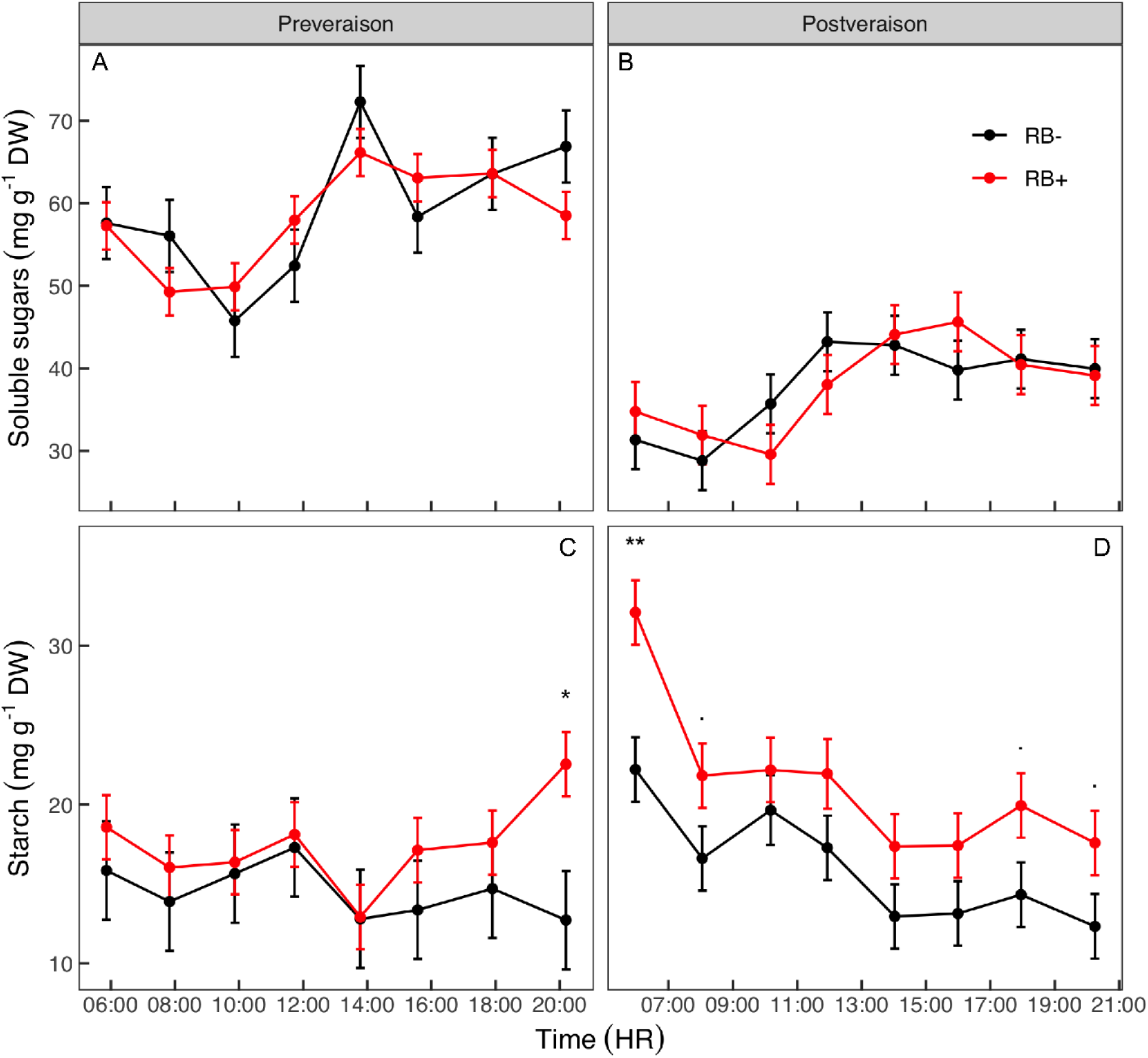
Responses of leaf soluble sugars (A, B) and starch (C, D) between healthy (RB-) and infected (RB+) vines over the course of the day on 23 July (preveraison; A, C) and 3 September (postveraison; B, D). Data are means ± one standard error (n = 3-7). ‘***,’ ‘**,’ ‘*,’ and ‘.’ represent statistically significant differences between means at *p* < 0.001, 0.01, 0.05, and 0.10, respectively.

Starch concentration in leaves, however, was consistently higher in RB+ leaves compared to RB-leaves across measurement times during both sampling dates (Figs. 4C-D). The leaf starch concentration was higher (ranging 14-91%) in RB+ vines during late afternoon (past1600 hr.) compared to RB-vines at preveraison, however, the difference was statistically significant only after sunset (2000 hr.). Average leaf starch concentration preveraison was 20% higher (*p* = 0.115) in RB+ vines (17.4 ± 0.9 mg g^-1^ DW) compared to RB-vines (14.5 ± 1.4 mg g^-1^ DW). Postveraison, the starch concentration in RB+ leaves was higher at all time points throughout the day (Fig. 4D). On average, leaf starch concentration was ∼ 32% higher (*p* = 0.029) in RB+ vines (21.3 ± 1.4 mg g^-1^ DW) compared to RB-vines (16.1 ± 1.4 mg g^-1^ DW). However, significant differences between individual means were only observed during early morning and late afternoon.

### Carbon export

Like *A*_net_, GRBV had a noteworthy impact on carbon export especially during late afternoon postveraison (Figs. 5B and 5D). Averaged across the afternoon (1300 to 1900 hr.) measurements, postveraison carbon export was ∼46% lower (*p* = 0.033) in RB+ leaves compared to RB-leaves. Average *A*_net_ for RB+ leaves (8.6 μmol m^-2^ s^-1^) was slightly higher than export rate (7.1 μmol m^-2^ s^-1^) during the afternoon period, indicating that carbon was not being exported as quickly as it was assimilated. Although there were no significant differences in carbon export between RB- and RB+ vines preveraison, afternoon export in RB+ trended lower compared to RB-.

**Figure 5.**
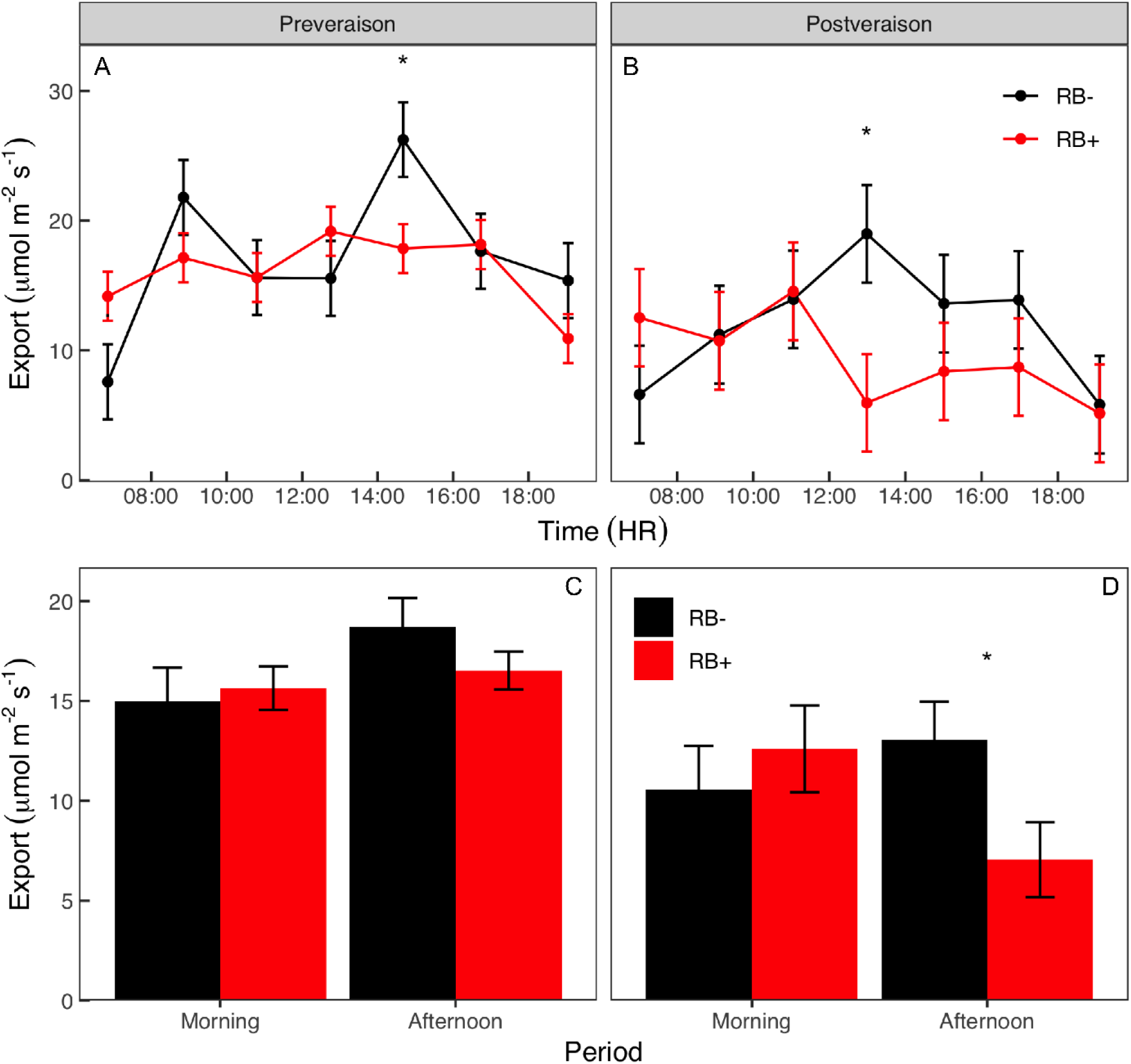
Response of leaf carbon export rate between healthy (RB-) and infected (RB+) vines over the course of the day (A, B) and averaged during morning and afternoon (C, D) on 23 July (preveraison; A, C) and 3 September (postveraison; B, D). Data are means ± one standard error (n = 3-7). ‘*’ represents statistically significant differences between means at *p* < 0.05.

### Chronology of disease symptoms

Over the course of the season, foliar starch accumulation in RB+ vines preceded the delayed ripening symptoms in the fruit, which in turn preceded the expression of disease symptoms in leaves (Fig. 6). The leaf starch concentration of RB+ vines was consistently, however not significantly, higher than that of RB-vines 14 days prior to veraison (Fig. 6A). Fruit ripening, indicated by sugar accumulation in berries, in RB+ vines did not start to lag behind RB-vines until 10 days postveraison, at which point sugar per berry was significantly lower in RB+ vines compared to RB-vines (Fig. 6B). Similarly, berry anthocyanin content was significantly lower in RB+ vines compared to RB-vines 10 days postveraison (Fig. 6C). Expression of red leaf symptoms (i.e., disease severity) increased significantly in RB+ vines following the delayed ripening symptoms in the fruit (Fig. 6D).

**Figure 6.**
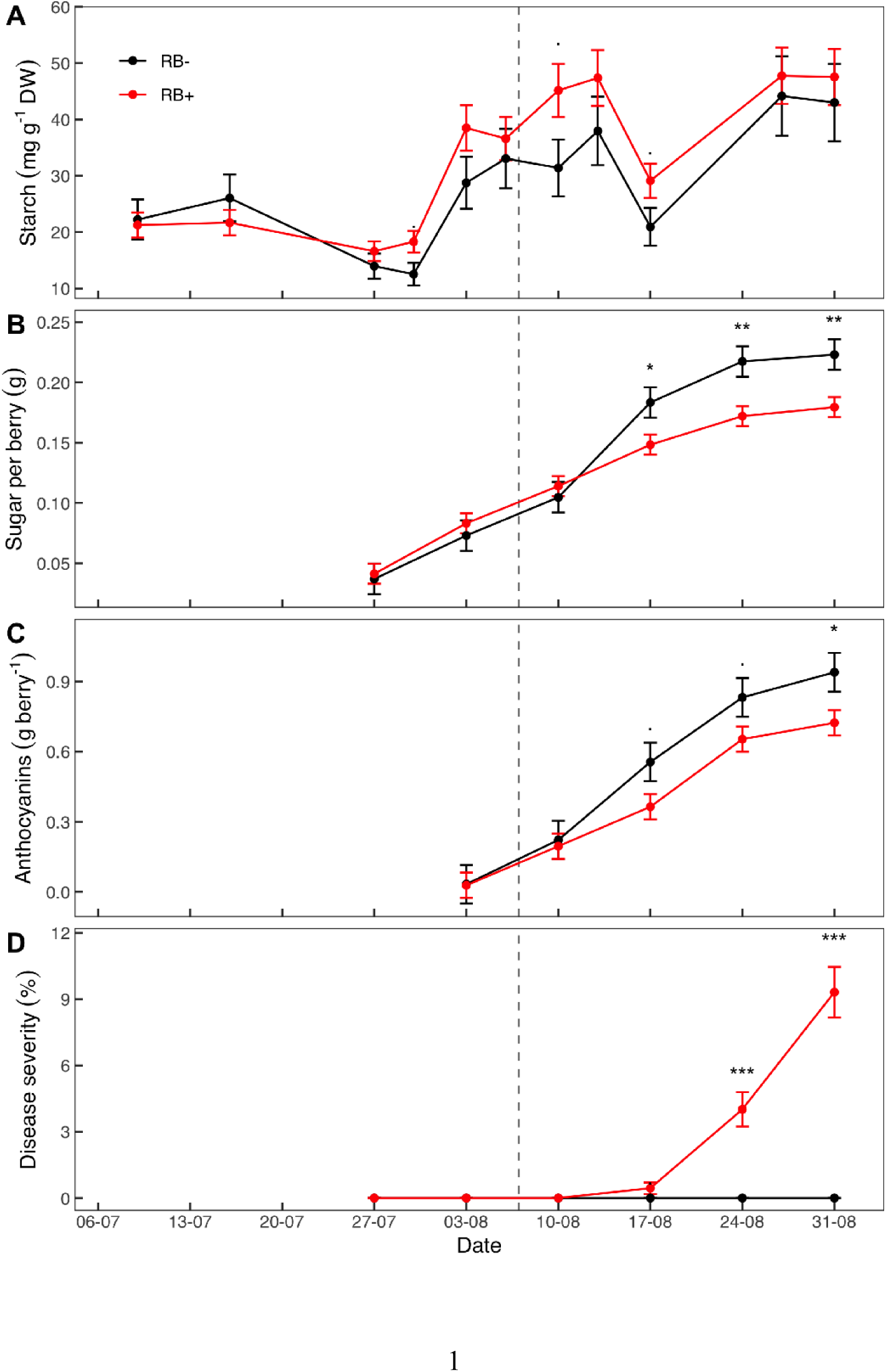
Responses of leaf starch concentration (A), berry sugar content (B), berry anthocyanin content (C), and disease severity (D) between healthy (RB-) and infected (RB+) vines over the course of the growing season. The vertical dashed line indicates the date of veraison. Data are means ± one standard error (n = 3-7). ‘***,’ ‘**,’ ‘*,’ and ‘.’ represent statistically significant differences between means at *p* < 0.001, 0.01, 0.05, and 0.10, respectively.

## Discussion

This study utilized seasonal and diurnal sampling strategies in combination with a mass balance approach to test the hypothesis that GRBV infection directly reduces leaf carbon export that results in a feedback inhibition of photosynthesis due to leaf starch accumulation. The results support this hypothesis and furthermore demonstrate that impacts of GRBV on leaf carbon metabolism vary seasonally and diurnally, with strongest adverse impacts occurring postveraison and in the afternoon, corresponding to the period of greatest light-utilization as well as highest water and heat stress. Finally, the results establish a chronology of disease symptom development that begins with the accumulation of starch in leaves preveraison, followed by reduced leaf carbon export, reduced *A*_net_, reduced sugar accumulation in berries, reduced anthocyanin biosynthesis in berry skins, and lastly increased anthocyanin biosynthesis in leaves. With this new understanding of starch accumulation as an earlier biomarker of infection, a chronology of GRBV symptom expression begins to emerge which may guide future detection and management of the virus.

It is well established that GRBV reduces *A*_net_ later in the growing season around and after veraison (Martinéz-Lüscher et al. 2019, Bowen et al. 2020, Levin and KC 2020). Reduced sugar translocation to fruit in RB+ vines was inferred by the coincidence of lower postveraison *A*_net_, elevated postveraison foliar NSC, and reduced sugar in the berries (Martinéz-Lüscher et al. 2019). However, this study is the first to directly quantify the impact of GRBV on carbon export. The data presented here demonstrate that the impact of GRBV on carbon metabolism is transient throughout the course of a day and may be most severe later in the day when environmental stress is the greatest. Additionally, this study provides further evidence that the impact of GRBV on carbon metabolism is driven by end-product inhibition of photosynthesis and mediated primarily by accumulation of starch in RB+ leaves. This accumulation of starch is the earliest detectable symptom in infected vines that appeared two weeks before veraison, well before a reduction in photosynthesis or appearance of red leaves were detected. This finding suggests a potential early detection biomarker of GRBV infection to facilitate management practices in vineyards.

### The impact of GRBV on carbon assimilation and export varies diurnally

The diurnal pattern of GRBV impact on gas exchange correlates with the degree of environmental stress as far as *A*_net_ and carbon export are reduced by a greater magnitude later in the afternoon when air temperature and VPD are higher. In addition, this is the same period of the day when Ψ_leaf_ reaches a daily minimum. Previous study of the effect of GRBV on grapevine water relations report that disease symptoms are exacerbated by water deficits (Levin and KC, 2020; Copp and Levin, 2021). In contrast, maintaining a high plant water status has been suggested as a way to improve *A*_net_ in leaves and sugar accumulation in the fruit of GRBV-infected leaves (Copp and Levin, 2021). Following the pressure-flow hypothesis of phloem function (Münch, 1927), Copp and Levin (2021) hypothesized that maintaining a higher water status in RB+ leaves would increase the gradient of water flow in the phloem, subsequently improving sugar translocation out of the leaf. Thus, the lower Ψ_leaf_ later in the day may further hinder phloem loading in GRBV-infected leaves and cause increased accumulation of NSC and subsequent feedback inhibition of photosynthesis. The mechanism, however, of inhibited translocation in the phloem in RB+ vines remains an open question but may be linked to physical blockages at the phloem sieve plates, as has been demonstrated in other plant pathosystems (Welker et al., 2022).

This study shows that *A*_net_ and carbon export rate are not impacted to the same extent by GRBV during the day, even postveraison when the impacts of the disease are assumed to be most severe (Levin and KC 2020). Specifically, the late afternoon reduction in *A*_net_ and carbon export has obvious consequences for the physiological impact of GRBV during the ripening period, but also significant implications for gas exchange sampling and study of GRBV-infected vines. Gas exchange is often measured in plants when irradiance is saturating and at steady-state to capture steady-state *A*_net_ and/or *g*_sw_ (e.g., at solar noon). However, the maximum impact of GRBV on *A*_net_ observed in this study may be mismatched with the windows for steady-state irradiance or *A*_net_. This field study was instigated, in small part, to evaluate whether the standard sampling window for gas exchange measurements (approx. 1100 –1300 hr.) failed to capture the full impact of GRBV on gas exchange. Though the present study does not contain a continuum of diurnal response of gas exchange to GRBV throughout the season, the data demonstrate that regular sampling of gas exchange may not always capture maximum differences between RB- and RB+ vines before 1200 hr. even two weeks postveraison.

### GRBV increases foliar starch, but not soluble sugars

The present study only found higher leaf starch in RB+ vines compared to RB-vines in contrast to other studies on foliar NSC in RB+ vines that reported elevated levels of soluble sugars (Wallis and Sudarshana, 2016; Martínez-Lüscher et al., 2019). The diurnal changes in soluble sugar concentration observed in our study followed the previously reported pattern of sucrose concentration in grapevine leaves increasing earlier during the day before saturating in the afternoon (Chaumont et al., 1994).

Beyond this point during the preveraison sampling date, when soluble sugars were saturated in both RB+ and RB-vines, there was an observed increase in starch concentration in RB+ vines whereas in RB-vines starch trended downward as the day progressed. Postveraison, however, the saturation in soluble sugars in RB-vines was achieved at least four hours prior (1200 hr.) to RB+ vines (1600 hr.). This different temporal dynamic may explain why other studies reported differences in soluble sugars between RB+ and RB-vines. Nevertheless, leaf starch trended downward for both RB+ and RB-vines through the course of the day postveraison except for a brief but significant increase past the leaf soluble sugar saturation point in RB+ vines at the end of the day. Thus, the excess leaf carbohydrate synthesized after this saturation point in RB+ vines was diverted into starch synthesis as previously shown in C_3_ plants (van Bel and Hafke, 2005). Likely this was because of blockage of the phloem due to a defense response (Lewis et al., 2022) and/or was reallocated for other metabolism such as the anthocyanin biosynthesis that gives rise to the characteristic red blotches for which the disease is named.

Starch is the primary carbohydrate of concern in the context of the end-product accumulation or feedback inhibition hypothesis governing the negative impact of GRBV on *A*_net_. The data herein align with previous description of the role of starch in end-product inhibition of photosynthesis across many plant species (Goldschmidt and Huber, 1992; van Bel and Hafke, 2005; Turgeon, 2010). Goldschmidt and Huber (1992) described how elevated starch concentration from girdling had the strongest relationship with inhibition of *A*_net_ across species and various carbohydrates. The concentrations of sucrose, fructose, and glucose in response to vascular blockage, such as girdling, vary across species and by invertase activity, but these carbohydrates are more transient forms than starch. Because chloroplasts are simultaneously the site of carbon assimilation and starch storage (Hummel et al., 2010), there seems to be a strong relationship between end-product regulation of photosynthesis and starch accumulation in GRBV-infected grapevines. Anatomical studies of chloroplasts of RB- and RB+ vines would confirm the role of starch in photosynthetic feedback inhibition in a GRBV context.

If accumulation of foliar starch due to inhibited carbon export subsequently reduces *A*_net_ in RB+ vines, then the starch signal could be detectable before other changes to carbon metabolism. In the measurements conducted over the course of the season, increases in foliar starch in RB+ vines relative to RB-vines was observed prior to veraison and was at least 2 weeks prior to the first observation of red leaf symptoms. This may be useful for guiding sampling related to the impacts of GRBV on vine physiology and carbon metabolism but has even greater potential to inform early detection of the disease using remote sensing techniques (Tanner et al., 2022). For example, hyperspectral imaging has been used for early detection of Huanglongbing (citrus greening) through changes to foliar NSC concentrations, which tend to be higher in leaves from trees with Huanglongbing (Weng et al., 2018). Thus, the elevated foliar starch levels shown here have the potential to serve as biomarkers to guide the generation and timing of hyperspectral imaging protocols for early identification of GRBV without intensive lab-based virus detection methods (DeShields and KC, 2023).

### GRBV-infected vines exhibit no sustained impairment of photochemical efficiency

The lack of differences in *F*_v_/*F*_m_ values between RB- and RB+ vines observed in this study suggests that there is no sustained impairment of photochemical efficiency due to GRBV infection. The values measured ranged from 0.79 – 0.81 and indicated a non-stressed status (Gallé and Flexas, 2010) even in RB+ vines. The lack of differences indicates that GRBV had no impact on the efficiency of photosystem II. Light adapted fluorescence (Φ_PSII_) was used to approximate photochemical efficiency during the day and was reduced in RB+ leaves relative to RB-leaves. This trend followed those of *A*_net_ and carbon export to the degree that Φ_PSII_ values in RB+ leaves were only significantly lower than those in RB-leaves in the afternoon during the postveraison sampling date. In tandem with transiently reduced *J* and *qP*, the reduction in Φ_PSII_ suggested that the quanta of light absorbed was more than that required for carbon assimilation (Gallé and Flexas, 2010). This transient reduction in Φ_PSII_, *J*, and *qP*, and a lack of reduction in *F*_v_/*F*_m_ in RB+ vines compared to RB-vines support the hypothesis of an end-product feedback inhibition of photosynthesis as opposed to sustained damage or alteration of photochemical efficiency (Pammenter et al., 1993).

Quantification of *NPQ* showed no response to virus status, which was curious considering the reduction in photochemical quenching in RB+ vines inferred from reductions in Φ_PSII_, *J*, and *qP*. *NPQ* calculation usually requires *F*_m_ values from dark-adapted leaves, but predawn *F*_m_ values were substituted here in the calculation of *NPQ*, so interpretation of *NPQ* values may be limited. In leaves with reduced *qP*, elevated *NPQ* values would be expected to indicate the initiation of processes like the xanthophyll cycle to protect the leaf from light damage (Gallé and Flexas, 2010). For RB+ vines, foliar anthocyanin synthesis may fulfil this role (Liakopoulos et al., 2006). In this study, however, all ChlF measurements were made on asymptomatic (i.e., green) leaf portions and thus the excess light energy would need to be preliminary dissipated in another manner. Additional leaf phytochemical analysis may explain the non-photochemical quenching capacity and photoprotective strategies of RB+ leaves both before and after anthocyanin biosynthesis.

### The expression of disease symptoms is temporally linked to carbon metabolism

The measurements conducted throughout the ripening period indicated that reduced anthocyanin biosynthesis in berries followed the reduced sugar accumulation in the berries in RB+ vines, but preceded anthocyanin biosynthesis in foliar tissues. While Martínez-Lüscher et al. (2019) first linked the reduction in anthocyanins to lower sugar in the fruit, the data in this study clearly show the temporal nature of this relationship. The appearance of red leaf symptoms follows the reduction in fruit metabolite accumulation that itself follows the increased accumulation of NSC in the leaves. This signal triggers foliar anthocyanin biosynthesis as excess solar radiation cannot be quenched through photochemistry in RB+ leaves and excess sugar is available as a substrate (Feild et al., 2001). The role of carbohydrates in foliar and fruit anthocyanin synthesis is well documented, as sucrose is both the signal and substrate for anthocyanin synthesis (Pirie and Mullins, 1976; Lecourieux et al., 2014). However, the coordination of these GRBV-mediated processes has not been reported previously.

Though the data presented here show how the appearance of these foliar and fruit symptoms are temporally linked, the onset and severity of disease symptoms is dependent on the response of carbon metabolism to the interaction of the virus and environmental factors. The association between severity of GRBV symptoms and stress suggest that foliar anthocyanin synthesis and accumulation of sugar and anthocyanins in the berries of RB+ vines relative to RB-vines could be accelerated or decelerated by water deficits or supplements, respectively (Levin and KC, 2020). For example, in this study, foliar reddening symptoms were first observed approximately 10 days postveraison at a berry TSS range of 16-17 °Brix. By contrast, Levin and KC (2020) first observed foliar reddening symptoms earlier with respect to both veraison (approximately 5 days pre- and 1 day postveraison) and TSS (6-7 and 12-13 °Brix; unpublished data), however that study investigated the impact of deficit irrigation on RB+ vines which may in part explain why symptoms manifested earlier. Indeed, the variability of disease symptom onset and severity across seasons and growing locations has been previously reported (Girardello et al., 2019; Levin and KC, 2020; Rumbaugh et al., 2021) and is one of the most challenging aspects of studying this plant pathosystem.

## Conclusion

This study elucidates the cascade of GRBV-mediated physiological symptoms beginning with impaired carbon export, followed by accumulation of foliar starch, causing a feedback inhibition of *A*_net_, and finally synthesis of foliar anthocyanin, coupled with reduction in sugar accumulation and anthocyanin synthesis in the fruit. This study also demonstrates that GRBV causes a considerable diurnal variation in the response of carbon assimilation and export. It also reveals the transient nature of the impact of GRBV on carbon metabolism. The impact of the virus and expression of symptoms are primarily caused by end-product feedback inhibition of photosynthesis and altered in severity and progression by stress (e.g., environmental and vine water stress). In addition to the broadened understanding of GRBV, this study provides insights into the development of “invisible” symptoms (e.g., starch accumulation) that could guide future hyperspectral detection of disease prior to the appearance of visible foliar symptoms.

## MATERIALS AND METHODS

### Vineyard site and plant material

The study was conducted in a 3.9 ha commercial block of *Vitis vinifera* L. cv. Pinot noir (clone 777) grafted on to 3309 Couderc (*V. riparia × V. rupestris*) rootstock. The vineyard site was in the Rogue valley AVA in southern Oregon (42°19’05.3” N, 122°56’18.5” W; 434 m above sea level). The vineyard was established in 2010 with a North-South row orientation, and 2.0 m × 1.5 m of row × vine spacing. The soil was comprised of both Medford and Gregory silty clay loams with less than 3% slope. Vines were trained on a vertically shoot positioned (VSP) trellis, and spur-pruned with bilateral cordons. The fruiting wire was located 1.0 m above the soil surface with two sets of foliage catch wires at approximately 1.3 m and 1.6 m above the soil surface. All vineyard management practices were conducted according to industry standards for the region.

### Determination of vine infection status and classification

Grapevines were sampled and tested for GRBV at commercial harvest of the 2019 growing season. Tissue samples consisted of four entire leaves (leaf blade + petiole), two from each cordon. Petioles were cut into ∼1 mm slices using a sterile razor blade, of which 100 mg per sample was transferred to microtubes for DNA extraction. DNA extractions were performed using a modified CTAB DNA extraction protocol (Richards et al., 1994). The diagnostic primers used in PCR amplification were CPfor/CPrev for the GRBV coat protein gene fragment, REPfor/REPrev for the GRBV replication-associated gene fragment, and 16Sfor/16Srev as a grapevine internal control for the 16S rDNA gene fragment (Krenz et al., 2014).

Disease symptom severity was quantified weekly from the onset of foliar reddening throughout the ripening period in 2020. Severity was estimated as the percentage of symptomatic leaves exhibiting any foliar reddening. Raw percentage data were converted into midpoint percentage values for analysis (Horsfall and Barratt, 1945). Grapevines that displayed characteristic symptoms of GRBD during the 2020 growing season, but tested negative for GRBV in 2019, were considered GRBV-infected and reclassified.

### Experimental design and measurement procedure

Diurnal measurements were conducted on two dates in 2020: 23 July (preveraison) and 3 September (postveraison). Measurements and samples were collected every two hours starting before dawn (0400-0500 hrs.) from previously identified grapevines of two virus statuses – healthy (RB-) or infected (RB+), for a total of 10 vines across 8 sampling times per date. At each sampling time, one leaf per vine was analyzed for gas exchange and chlorophyll fluorescence, then excised for measurement of Ψ_leaf_, and then stored at approximately 5 °C for later lab analysis of NSCs (details below). Due to the appearance of symptoms in data vines previously classified as RB-, new RB-vines were identified prior to the postveraison data collection period. Vines previously classified as RB-in the preveraison period were subsequently reclassified as RB+ prior to analyses.

Apart from the pre- and postveraison diurnal measurements, seasonal measurements were also conducted on the same vines in 2020. At regular intervals throughout the growing season, the same data were collected as in diurnal measurements described above, but at a single time point in the day (solar noon). In addition, berry samples were collected for maturity analyses (details below), and vines were visually rated for disease severity as described in Levin and KC (2020).

### Environmental conditions

Air temperature and relative humidity data were accessed from a nearby weather station (MDFO, AgriMet, United States Bureau of Reclamation) located approximately 1.8 km from the study site (42.3311°N, 122.9377°W). Vapor pressure deficit (VPD) was calculated from air temperature and relative humidity. Photosynthetically active radiation (PAR) data were collected using a photosynthetic photon flux density (PPFD) sensor attached to a portable photosynthesis system (LI-6400XT, LI-COR Biosciences, Lincoln, NE) and determined at the time of each gas exchange measurement.

### Leaf gas exchange, chlorophyll fluorescence, and water status

Leaf gas exchange was measured as *A*_net_ and *g*_sw_ at each sampling point using the portable photosynthesis system described above. Two fully expanded, sunlit (when sun was present) leaves per vine were used for measurement. Relative humidity, temperature, and PPFD were set in the leaf chamber to match ambient conditions. Flow rate was set at 400 µmol s^-1^, chamber CO_2_ concentration was set in the reference cell at 400 µmol mol^-1^. Measurements were made once *A*_net_ and *g*_sw_ had stabilized, which took 60-90 s per leaf. Infrared gas analyzers were matched every 20 min during measurement periods.

Chlorophyll fluorescence was measured simultaneously with gas exchange using a pulse-amplitude modulated (PAM) fluorometer (6400-40, LI-COR Biosciences, Lincoln, NE). *F*_v_/*F*_m_ was determined only for dark-acclimated leaves at the predawn time points on each measurement date. Subsequent analyses showed no differences between healthy and infected vines at each measurement date, thus predawn *F*_m_ values were used in the calculation of light-acclimated parameters. For light-acclimated leaves, the quantum yield of photosystem II (Φ_PSII_), linear electron transport rate (*J*), proportion of open PSII reaction centers (*qP*), and non-photochemical quenching (NPQ) were calculated. Φ_PSII_ was calculated using the following equation in which *F*_m_’ is maximal fluorescence of the light-acclimated leaf and *F*_s_ is the steady state fluorescence of the light-adapted leaf:

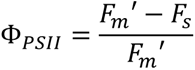

Non-photochemical quenching parameter (NPQ) was calculated using the following equation in which *F*_m_ was substituted from the pre-dawn measurements averaged per GRBV status:

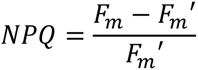

linear electron transport rate (*J*) was calculated using the following equation in which *f* is the fraction of absorbed quanta used by photosystem II (assumed 0.5), PPFD is photosynthetic photon flux density at the time of measurement, and *α_leaf_*is leaf light absorptance recorded by the system:

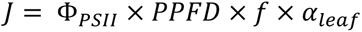

Ψ_leaf_ was determined using a pressure chamber (Model 615, PMS Instruments, Albany, OR) according to Levin (2019) for the same fully expanded, sunlit leaves (except prior to sunrise) as used for gas exchange and fluorescence measurements.

### Extraction and quantification of nonstructural carbohydrates

Leaves that were used in the gas exchange, fluorescence, and Ψ_leaf_ measurements were removed immediately after Ψ_leaf_ determination, placed in a paper bag, returned to the laboratory, and microwaved for 90 s at 600 W to halt metabolic activity (Landhausser et al., 2018). Samples were stored at -20 °C until oven-dried at 70 °C for 48 hr. Dried leaves were ground using a Mini Wiley Mill (Thomas Scientific, Swedesboro, NJ) and passed through a 40-mesh sieve.

Nonstructural carbohydrates were extracted as described in Chow and Landhäusser (2004) and Landhausser et al. (2018), with some modifications. Twenty-five mg of ground leaf tissue was mixed with 1 mL of 80% ethanol, vortex for 10 s, and incubated at 90 °C for 10 min. The ground leaf tissue suspension was then centrifuged at 13,000 x g for 1 min and 1 mL of supernatant was removed for soluble carbohydrate quantification. The remaining starch pellet was washed twice more, using the same volume of ethanol and all supernatant was discarded. After leaving the starch pellet to airdry for approximately 18 hours, the pellet was resuspended in a 600 U/mL α-amylase solution and incubated at 85 °C for 30 min and then centrifuged at 13,000 × *g* for 1 min. One hundred µL of supernatant was then mixed with 500 µL of 12 U/mL amyloglucosidase (AMG) solution and incubated at 55 °C for 30 min. The mixture was brought to room temperature before quantifying the glucose hydrolysate.

Glucose quantifications were performed with using an anthrone-sulfuric acid method at 620 nm, in accordance to Leyva et al. (2008). Dilutions of a 180 mg L^-1^ glucose solution were used as a standard curve. Five mg of potato starch was used as a starch control for determining the starch digestion efficacy (SDE). Samples and glucose controls were analyzed in technical duplicates and starch controls were analyzed in biological duplicates. Glucose and starch were calculated as mg/g of sample dry weight (mg/g DW) using the following formulae:

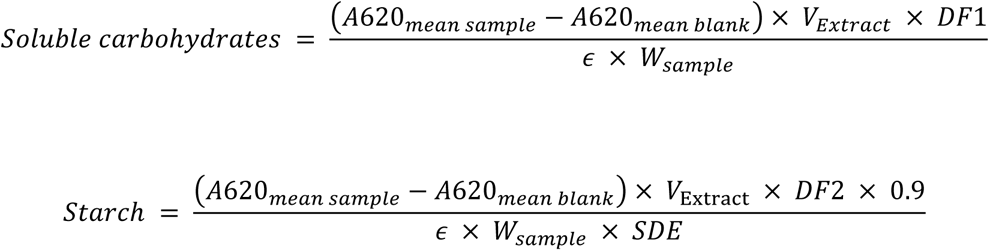

where A620_mean sample_ is the mean absorbance of the unknown sample duplicates; A620_mean blank_ is the mean absorbance of the reagent blank duplicates; V_extract_ is the volume of the extract expressed as L made with either ethanol or α-amylase solution; DF1 is the dilution factor of the soluble carbohydrate sample; DF2 is the dilution factor of the glucose hydrolysate sample multiplied by six or the dilution factor for the dilution of α-amylase solution in amyloglucosidase solution; 0.9 is the weight conversion factor of starch to glucose; ɛ is the absorption coefficient of the glucose standard; W_sample_ is the measured weight expressed as grams of dry tissue sample in the tube; and SDE is calculated using the starch formula using a W_sample_ of 0.005 g.

### Estimation of leaf carbon export

Leaf carbon export was estimated using a mass balance approach as outlined in Dayer et al. (2021) and Gersony et al. (2020) using the following formula:

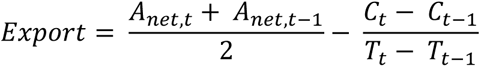

where export is the difference between average *A*_net_ and the change in carbon concentration (*C*) between two time points (*t* and *t-1*). *C* is the total quantity of NSCs per unit leaf area converted into molar units compatible with *A*_net_ using the molecular weight of glucose. The export rate is expressed as μmol C m^-2^ s^-1^ and is plotted in figures at the midpoint between *t* and *t-1*.

### Berry sugar content and skin anthocyanins

Sugar per berry and anthocyanin content were quantified at six and five time points, respectively, throughout ripening, starting at approximately 10 days prior to the onset of ripening (veraison). For each replicate vine at each time point, 20 berries were sampled, weighed, and 10 berries were subsampled to be stored at -20°C for later anthocyanin analysis. The remaining 10 berries were crushed, and the juice was centrifuged at 15,000 × *g* for 5 min before being analyzed for total soluble solids (TSS) using a digital refractometer (AR200, Reichert Analytical Instruments, Depew, NY). Sugar per berry was estimated as the product of berry mass and TSS as in Krasnow et al. (2009). The 10-berry subsamples were peeled, the skins then dried, and extracted in 70% acetone for 24 hr. at 100 rpm on an orbital shaker (VWR, Radnor, PA). Acetone was removed from skin extracts by vacuum distillation (Syncore Analyst Polyvap, BUCHI Corporation, New Castle, DE). Anthocyanins were then quantified from the distilled extracts using the Harbertson-Adams assay (Harbertson et al., 2002; Heredia et al., 2006).

### Statistical analyses

Statistical analyses were conducted using R software for statistical computing (v. 4.0.3; www.R-project.org). For diurnal and seasonal datasets, two-way ANOVAs were conducted on data fit with linear models corresponding to a split-plot design in which viral infection status was the main plot factor and sample time (or date) was the sub-plot factor. Models were fit with the *lmer()* function from the *lmerTest* package (Kuznetsova et al., 2017) to account for the imbalance. For diurnal data, pre- and postveraison datasets were analyzed separately, due to the spread of the viral disease as previously mentioned. At the end of the season, n = 3 and 7 for RB- and RB+ preveraison, respectively, while n = 5 for both statuses postveraison. Means were estimated and compared using the *emmeans* package (v.1.6.3; Lenth, 2022). Figures were generated with the *ggplot2* package (v.3.3.5; Wickham, 2016).

## Acknowledgements and Funding

The authors thank Grestoni Vineyards, LLC for field plot maintenance and for providing the study site, and the Rogue Valley Winegrowers Association for continued support of the viticulture research program at SOREC. This work was financially supported in part by the Oregon Wine Research Institute’s Undergraduate Scholars program and the USDA-NIFA-SCRI (grant number 2019-51181-30020).

## Author Contributions

A.K.C. and A.D.L. obtained funding for the research; C.R.C. and A.D.L. designed the research; C.R.C., J.B.D., C.K., R.C., and A.D.L. collected the field data; J.B.D. created the nonstructural carbohydrate protocol and assisted with troubleshooting; C.R.C., J.B.D., and C.K. performed the nonstructural carbohydrate measurements; C.R.C. and M.S. performed the berry measurements; C.R.C., J.B.D., and A.D.L. analyzed the data; C.R.C. wrote the first draft of the paper; C.R.C., J.B.D., S.K., and A.D.L. edited the final draft.

